# Pharmacological stimulation of infralimbic cortex after fear conditioning facilitates subsequent fear extinction

**DOI:** 10.1101/2024.03.23.586410

**Authors:** Hugo Bayer, James E. Hassell, Cecily R. Oleksiak, Gabriela M. Garcia, Hollis L. Vaughan, Vitor A. L. Juliano, Stephen Maren

**Affiliations:** Department of Psychological and Brain Sciences, Texas A&M University, College Station; Department of Pharmacology, São Paulo University, São Paulo

## Abstract

The infralimbic (IL) division of the medial prefrontal cortex (mPFC) is a crucial site for extinction of conditioned fear memories in rodents. Recent work suggests that neuronal plasticity in the IL that occurs during (or soon after) fear conditioning enables subsequent IL-dependent extinction learning. We therefore hypothesized that pharmacological activation of the IL after fear conditioning would promote the extinction of conditioned fear. To test this hypothesis, we characterized the effects of post-conditioning infusions of the GABA_A_ receptor antagonist, picrotoxin, into the IL on extinction of auditory conditioned freezing in male and female rats. In four experiments, we found that picrotoxin injections performed immediately, 24 hours, or 13 days after fear conditioning reduced conditioned freezing to the auditory conditioned stimulus (CS) during both extinction training and extinction retrieval; this effect was observed up to two weeks after picrotoxin infusions. Interestingly, inhibiting protein synthesis inhibition in the IL immediately after fear conditioning prevented the inhibition of freezing by picrotoxin injected 24 hours later. Our data suggest that the IL encodes an inhibitory memory during the consolidation of fear conditioning that is necessary for future fear suppression.

## INTRODUCTION

Animals rely on fear memories for survival. However, it is equally important for animals to suppress fear after danger has passed. Impairments in the extinction of fear memories is a key feature of post-traumatic stress disorder (PTSD) [1,2], but the precise reasons for why some memories are resistant to extinction are unclear. Several studies in humans and rodents show that the medial prefrontal cortex is crucial for extinction after Pavlovian fear conditioning [3-6]. In rodents, the infralimbic cortex (IL) is essential for the extinction of fear memories [3, 4, 7]. Pharmacological manipulations of the IL during extinction training [4, 5, 8, 9] or during the post-extinction consolidation period [8, 10, 11] influence the acquisition and retrieval of extinction memories.

In addition to its role in extinction learning, recent data suggest that the IL might have a role in other inhibitory learning processes. For example, IL inactivation reduces latent inhibition of fear, whereas stimulation facilitates extinction of appetitive conditioned responses [12, 13]. This suggests that the IL participates in many forms of inhibition, including inhibitory memories that might be encoded during fear conditioning itself. Interestingly, previous work by the first author has shown that pharmacological inactivation or protein synthesis inhibition in the IL immediately after contextual fear conditioning impairs subsequent extinction of fear [14]. Similarly, it was also found that silencing the IL several hours after fear conditioning leads to later extinction impairments [15]. This raises the intriguing possibility that the IL is engaged during (or soon after) Pavlovian fear conditioning to encode an inhibitory memory that supports later extinction of the excitatory fear memory. Consistent with this, there is evidence that aversive avoidance learning increases IL c-fos expression [16] and IL inactivation impairs the consolidation of avoidance [17, 18]. Additionally, Pavlovian fear conditioning increases c-fos expression in the IL [19,20]. Post-conditioning activation of the IL might represent the encoding of inhibitory engrams in the IL that are required for fear extinction.

Several studies have shown that pharmacological activation of the IL during extinction training facilitates long-term extinction memory [4, 5, 8, 9]. We hypothesized that pharmacological activation of the IL Here we characterized the effects of post-conditioning IL pharmacological stimulation and protein synthesis inhibition on subsequent extinction learning.

## MATERIALS AND METHODS

### Subjects

In this study we used 133 male and female Long-Evans rats obtained from Envigo (200-225g on arrival). Rats were individually housed in cages within a temperature- and humidity-controlled vivarium and kept on a 14:10 light:dark cycle (lights on at 7 AM). Experiments were performed during the light phase of the cycle. Rats had *ad libitum* access to standard rodent chow and water. Before behavioral testing, rats were handled for 5 days (∼30 sec/rat/day). All experimental procedures were conducted in accordance with the US National Institutes of Health (NIH) Guide for the Care and Use of Laboratory Animals and were approved by the Texas A&M University Institutional Animals Care and Use Committee (IACUC).

### Surgery

Rats were anesthetized with isoflurane (5% for induction and 1-2% for maintenance) and secured in a stereotaxic apparatus (Kopf Instruments, Tujunga, CA) for IL cannulation. Animals received a 10 mg/ml/kg injection of Rimadyl (Zoetis), the incision area on the scalp was shaved, povidone-iodine pads were used for antisepsis, ophthalmic ointment was applied to the eyes, and lidocaine was injected subcutaneously beneath the scalp. An incision was made using a scalpel blade and the scalp was retracted using hemostats. The skull was leveled to place bregma and lambda in the same horizontal plane, and holes were drilled in the skull for placement of the cannula and two jeweler’s screws, which were affixed near the lambdoid suture.

Stainless steel guide cannulae (26 gauge, 9-mm from base of the pedestal; Plastics One or RWD) were implanted using the following coordinates [mm relative to either bregma (AP and ML) or skull surface (DV)]: AP = + 2.65, ML = ± 3.2, DV = -6.3 to -6.5 at a 30° angle. After the cannulae were lowered, dental cement was applied to the skull surface to secure the immediately after fear conditioning would also facilitate the subsequent extinction of fear memory.

cannulae to the skull. After surgeries, topical antibiotic (Triple Antibiotic Plus, G&W Laboratories) was applied to the skin surrounding the headcap. Dummy cannulae (33 gauge, extending 1 mm beyond the cannula tip) were inserted into the guide cannulae to prevent clogging. Animals had at least one week for recovery between surgery and behavioral testing.

### Drugs and infusion procedures

For drug infusions, animals were transported to the infusion room and stainless-steel injectors (33 gauge) connected to Hamilton syringes mounted on infusion pumps were inserted into the guides. Picrotoxin (PIC 200 ng/μL) was dissolved in 0.5% ethanol-saline and was used to stimulate the IL [21]. Brain-derived neurotrophic factor (BDNF; 2.5 μg/μl) was dissolved in saline and used to promote plasticity in the IL [22, 23]. Anisomycin (ANISO; 100 μg/μl) was dissolved in 5 N HCl and PBS (pH was corrected using NaOH) and was used to inhibit protein synthesis in the IL [14, 24, 25]. Injections (0.15 μl/min; 0.3 μl total) lasted two minutes and injectors were left in place for one more minute to allow diffusion.

### Behavioral apparatus and procedures

All experiments were conducted in 16 standard rodent conditioning chambers (30 × 24 × 21 cm; Med Associates, St Albans, VT), housed inside sound-attenuating cabinets. The chambers had two aluminum sidewalls, a Plexiglas rear, ceiling, and door, and a grid floor. The grid floor consisted of 19 stainless steel bars that were connected to a shock source, and a solid-state grid scrambler for foot shock delivery (Med Associates). Each conditioning chamber was equipped with a loudspeaker for delivering auditory stimuli, a ventilation fan and a light. Locomotor activity was transduced into an electrical signal by a load cell under the floor of the chamber to automatically measure freezing. Approximately one week after surgeries, rats underwent auditory fear conditioning in context A (chamber lights on, fans off, cabinet doors closed, 3% acetic acid, white transport boxes), which consisted of five tone-footshock pairings after a 3 min baseline period. The tones (CS) were 10 s, 80 dB, 2 kHz; the shocks (US) were 1 mA, 2 s, the intertrial interval (ITI) was 70 s long and animals remained in the chamber for an additional minute after the last pairing. Context exposure consisted of a 10 min stimulus-free period in the extinction context (context B; transport boxes with bedding). Extinction and extinction retrieval consisted of a 3-min stimulus-free baseline period followed by 45 CS-alone presentations (ITI = 40 s) followed by a 3 min post-trial period.

### Histological procedures

Rats were overdosed with sodium pentobarbital (100 mg/kg) and perfused transcardially with 0.1M PBS followed by 10% formalin. Brains were extracted from the skull and post-fix ed in a 10% formalin solution for 24h followed by a 30% sucrose solution where they remained for a minimum of 48 h. After the brains were fixed, coronal sections (30μm thickness) were made on a cryo-stat (−20°C), mounted on subbed microscope slides, and stained with thionin (0.25%) to visualize cannula placements. Only rats with bilateral IL cannula placements were included in the analysis.

### Statistical analysis

Data were analyzed with conventional parametric statistics (Prism 9.0; GraphPad Software). Freezing data for the pre-CS baseline (BL) period and for each trial (CS + ITI averages) are shown for conditioning and extinction sessions. Baseline freezing levels for the extinction retrieval sessions was minimal and is not shown. When appropriate, Fisher’s LSD post hoc tests were used for pairwise means comparisons. Group sizes were based on previous work from the laboratory. No significant sex differences were observed, so animals from both sexes were collapsed for all analyses (but are distinguished in bar graphs).

## RESULTS

### Experiment 1: Pharmacological stimulation of the infralimbic cortex immediately after fear conditioning suppresses freezing and facilitates extinction retrieval

To determine whether and how IL activation during fear memory consolidation affects fear retrieval and extinction learning, we infused animals with either VEH or PIC immediately after auditory fear conditioning. As shown in Figure 1B (left panel), all groups chamber lights off, fans on, cabinet doors open, 70% ethanol, black floors covering the grid floor, and black showed similar levels of freezing during the fear conditioning session and showed systematic increases in freezing across the session. This was confirmed in a two-way ANOVA, which revealed a main effect of conditioning trial (*F*_*5, 100*_ *=* 21.0, *p* < 0.01), but no other main effects or interactions (*F*s < 0.31, *p*s > 0.66).

**Figure 1.**
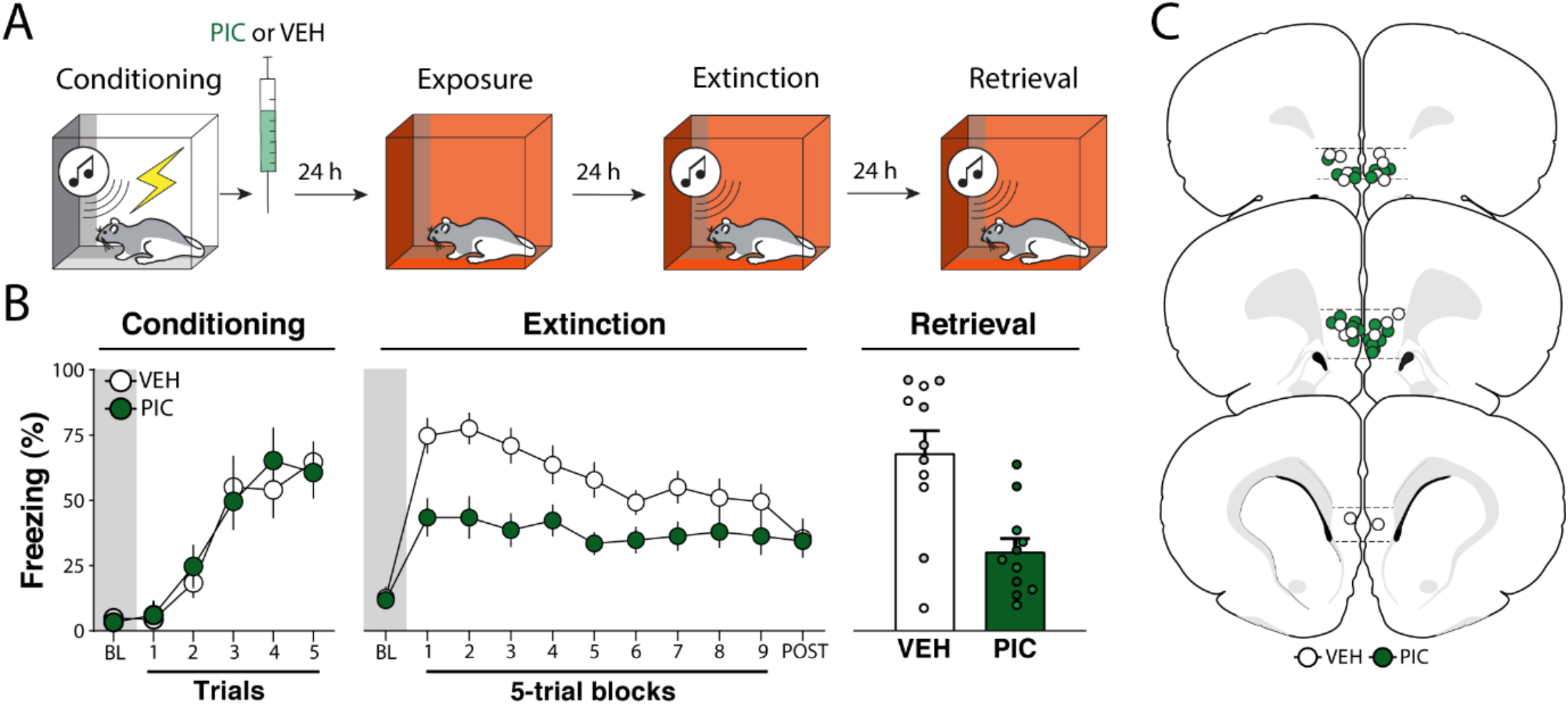
Experiment 1: Pharmacological stimulation of the infralimbic cortex immediately after fear conditioning suppresses freezing and facilitates extinction retrieval. **(A)** Behavioral schematic. **(B)** Freezing behavior during conditioning, extinction and retrieval are shown from left to right. During conditioning, both groups acquired conditioned freezing similarly. During extinction, picrotoxin (PIC) infusions in the infralimbic (IL) cortex led to a reduction in freezing across the session. During extinction retrieval, PIC-treated animals exhibited less freezing than vehicle (VEH)-treated animals. Retrieval data from male animals are shown in grey and dark green circles and squares, white and light green circles and squares represent data from female animals. **(C)** Schematic showing cannula placements in IL. Data are expressed as means ± standard error of the mean (s.e.m.). [PIC (*n* = 11); VEH (*n* = 11)], BL = baseline, PIC = picrotoxin, VEH = vehicle.

Two days after fear conditioning, extinction was performed. As shown in Figure 1B (middle panel), there were no group difference in baseline freezing (*t*_18_ = 0.190, *p =* 0.850). However, once the CS was delivered the PIC group exhibited lower freezing than controls throughout the extinction session. A two-way ANOVA performed for the entire session revealed main effects of trial (*F*_*9, 180*_ *=* 6.76, *p* < 0.01) and drug treatment (*F*_*1, 20*_ = 8.35, *p* < 0.01), with an interaction between the factors (*F*_*9, 180*_ *=* 2.75, *p* < 0.01). Although PIC reduced freezing during the first block of extinction trials, it did not affect freezing during the first CS (Figure S1). This suggests that PIC affected the extinction, rather than retrieval, of the fear memory. Despite showing lower levels of freezing at the outset of extinction training, VEH- and PIC treated rats came to exhibit similar levels of freezing by the end of the extinction session.

One day after extinction, animals were returned to the extinction context and tested for extinction retrieval. As shown in Figure 1B (right panel), the PIC group showed lower levels of freezing than controls during the 5-trial retrieval test (*t*_20_ = 3.74, *p* < 0.01). Hence, PIC treatment immediately after fear conditioning not only decreased freezing during the extinction session, but also facilitated extinction retrieval the following day.

### Experiment 2: Pharmacological stimulation of the infralimbic cortex 24 hours after fear conditioning suppresses freezing and facilitates extinction retrieval

Previous work has shown that BDNF infusions into the IL up to 24 hours after fear conditioning can cause a reduction in conditioned freezing similar to what we observed with immediate post-conditioning infusions of PIC [23]. Here we explored whether PIC infusions into the IL 24 hours after fear conditioning would also suppress freezing; we included a BDNF group as a positive control. As shown in Figure 2B (left panel), all groups increased freezing similarly across the conditioning session. This was confirmed by a two-way ANOVA, which revealed a main effect of conditioning trial (*F*_*5, 115*_ *=* 36.3, *p* < 0.01).

**Figure 2.**
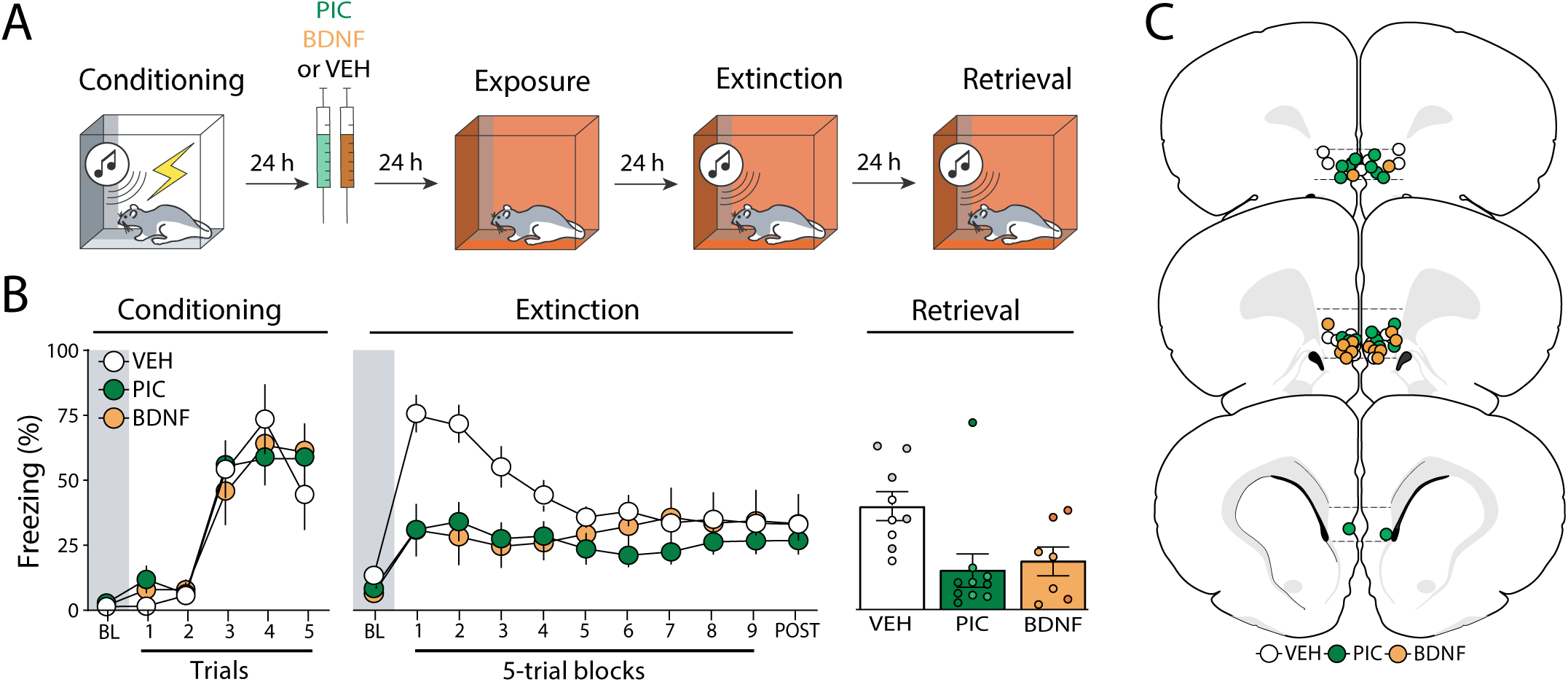
Experiment 2: Pharmacological stimulation of the infralimbic cortex 24 hours after fear conditioning suppresses freezing and facilitates extinction retrieval. **(A)** Behavioral schematic. **(B)** Freezing behavior during conditioning, extinction and retrieval are shown from left to right. During conditioning, all groups acquired conditioned freezing similarly. During extinction, intra-infralimbic (IL) cortical picrotoxin (PIC) or brain derived neurotropic factor (BDNF) infusions 24 h after fear conditioning reduced freezing behavior compared to vehicle (VEH)-treated animals. During the extinction retrieval test, PIC- and BDNF-treated animals also showed lower levels of freezing, but only the PIC significantly differed from controls. Retrieval data from male animals are shown in grey, dark green, and dark orange circles, while white, light green, and light orange circles represent data from female animals. **(C)** Schematic showing cannula placements in IL. Data are expressed as means ± standard error of the mean (s.e.m.) [VEH (n = 9); PIC (n = 10); BDNF (n = 7)], BDNF = brain-derived neurotrophic factor, BL = baseline, PIC = picrotoxin, VEH = vehicle.

Twenty-four hours after conditioning, animals received intra-IL infusions of VEH, PIC or BDNF and underwent extinction 48 hours later. As shown in Figure 2B (middle panel), there were no significant differences between groups (*F*_2,23_ = 1.55; *p* = 0.23) during the baseline period. However, animals that received infusions of PIC or BDNF exhibited lower freezing levels than VEH-treated animals once extinction trials were delivered. PIC two-way repeated measures ANOVA revealed significant main effects of trial (*F*_*9, 207*_ *=* 4.05, *p* < 0.01), treatment (*F*_*2,23*_ *= 3*.*51, p* = 0.05), and a significant interaction between these factors (*F*_*18,127*_ *=* 3.47, *p* < 0.01) on freezing behavior during the extinction session. Similar to the first experiment, PIC did not affect freezing during the first CS (Figure S1) and all rats reached similar levels of freezing by the end of the extinction session.

On the following day, animals were subjected to an extinction retrieval test. As shown in Figure 2B (right panel), both drug-treated groups showed lower levels of freezing than the control group. A one-way ANOVA revealed a significant effect of treatment (*F*_*2,23*_ = 5.28, *p* = 0.01) and post*-hoc* comparisons showed that both drug-treated groups had lower levels of freezing than the VEH group (*p*s < 0.03). Hence, PIC or BDNF infusions made 24 hours after conditioning (far outside of the classical consolidation time window) reduce the expression of conditional freezing during extinction and retrieval.

### Experiment 3: The effects of picrotoxin infusions in the infralimbic cortex are long-lasting

In the previous experiments, the interval between picrotoxin treatment and extinction training was never longer than two days. Therefore, it is possible that the disruption in freezing we observed were caused by an acute effect of the picrotoxin treatment. To investigate this possibility, we explored whether PIC infusions into the IL either 24 h or 13 days after fear conditioning would suppress freezing and facilitate extinction retrieval (Figure 3A).

**Figure 3.**
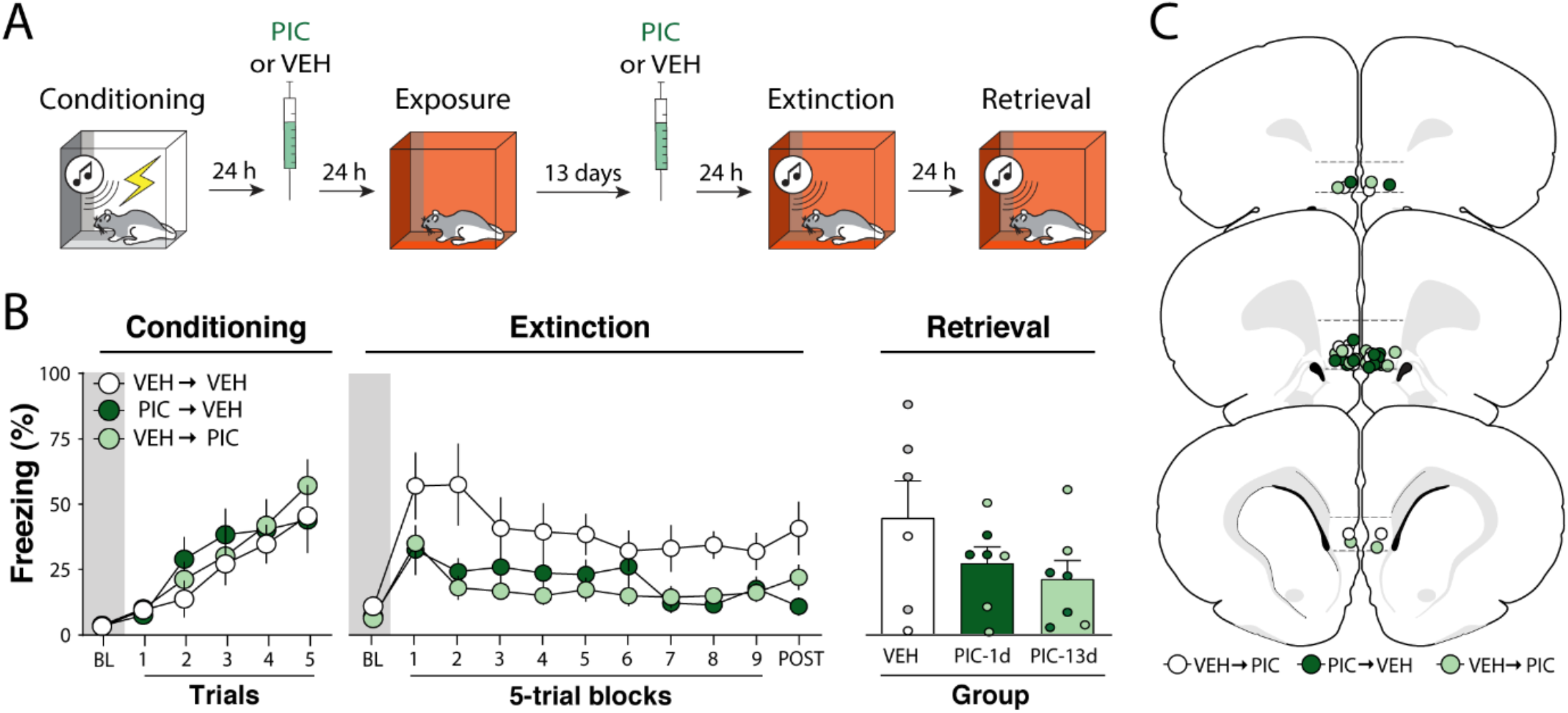
Experiment 3: The effects of picrotoxin infusions in the infralimbic cortex are long-lasting. **(A)** Behavioral schematic. **(B)** Freezing behavior during conditioning, extinction and retrieval are shown from left to right. During conditioning, all groups acquired freezing similarly. One and thirteen days after conditioning, animals received intrainfralimbic (IL) cortical infusions of picrotoxin (PIC) or vehicle (VEH). One day after the second intra-IL injection, animals underwent an extinction session; extinction retrieval was performed the following day. During extinction, animals that received intra-IL PIC 24 h or 13 d after fear conditioning showed lower levels of freezing. There were no group differences during the extinction retrieval test. Retrieval data from male animals are shown in grey and dark green circles, while white and light green circles represent data from female animals. **(C)** Schematic showing cannula placements in IL. Data are expressed as means ± standard error of the mean (s.e.m.) [VEH-VEH (*n* = 6); VEH-PIC (*n* = 7); PIC-VEH (*n* = 6)]; BL = baseline, PIC= picrotoxin, VEH = vehicle.

As shown in Figure 3B (left panel), all groups acquired conditioned freezing similarly during the conditioning session. An ANOVA revealed a main effect of trial (*F*_*5, 102*_ = 14.0, *p* < 0.01), but no other main effects or interactions (*F*s < 1.10, *p*s > 0.50). After receiving VEH or PIC infusions 1 or 13 days after conditioning, animals underwent extinction training 14 days after conditioning. As shown in Figure 3B (middle panel), baseline freezing before CS onset was similar in all groups (*F*_2,17_ = 1.09, *p* = 0.36). However, once the extinction session commenced, PIC infusions reduced freezing. This was confirmed by a two-way ANOVA, which revealed significant main effects of trial (*F*_*9,170*_ = 2.60, *p* < 0.01) and drug treatment (*F*_*2, 170*_ = 28.5, *p* < 0.01); the interaction of these factors was not significant (*F*_*18,170*_ = 0.55, *p* = 0.93). Furthermore, *post hoc* comparisons revealed that PIC the two PIC groups had lower freezing when compared to the control group (*p*s < 0.01) but were not significantly different from each other (*p* = 0.73). Similar to the first experiment, PIC did not affect freezing during the first CS (Figure S1).

One day after extinction, animals underwent a retrieval test. As shown in Figure 3B (right panel), no differences were observed during the test (*F*_2,18_ = 1.30, *p* = 0.30). Unlike the first two experiments, the PIC-induced suppression of freezing observed during the extinction session did not result in a significant enhancement of extinction retrieval the next day. Overall, these results reveal that intra-IL PIC infusion performed 24 hours after conditioning produce a long-lasting (13 day) decrement in subsequent conditional freezing during extinction training. Moreover, fear memories remain susceptible to intra-IL PIC treatment for at least 13 days after conditioning. However, in neither case did intra-IL PIC infusions facilitate extinction retrieval when extinction was conducted 48 hours after PIC infusions. These results suggest that PIC must be delivered within 24 hours of fear conditioning to enhance long-term extinction memories.

### Experiment 4: Protein synthesis inhibition in the infralimbic cortex during fear memory consolidation blocks the effects of picrotoxin

The previous experiment suggests that pharmacological stimulation of the IL outside of the consolidation time-window can reduce fear expression during extinction. However, a recent study suggests that protein synthesis inhibition in the IL during consolidation can impair posterior extinction of fear [14]. Therefore, we performed an experiment combining protein synthesis inhibition immediately after learning and pharmacological stimulation 24 h after learning (Figure 4A).

**Figure 4.**
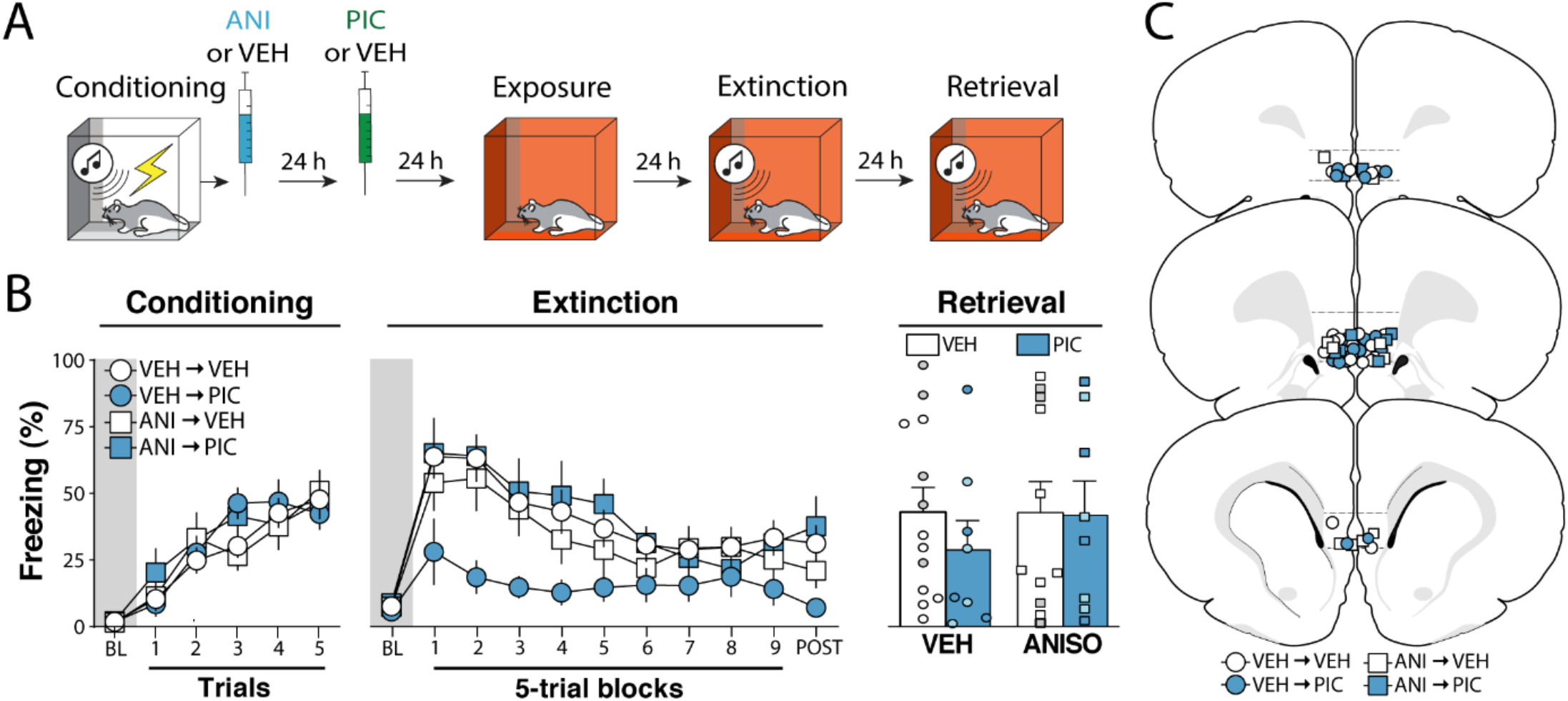
Experiment 4: Protein synthesis inhibition in the infralimbic cortex during fear memory consolidation blocks the effects of picrotoxin. **(A)** Behavioral schematic. **(B)** Freezing behavior during conditioning, extinction and retrieval are shown from left to right. All groups acquired conditioned fear similarly during the conditioning session and received intra-infralimbic (IL) cortical infusions of anisomycin (ANISO) or vehicle (VEH) immediately after. One day later animals were treated with either picrotoxin (PIC) or VEH and returned to the home cages for another day. Immediate post-conditioning ANISO infusions prevented the impairment in conditioned freezing observed in PIC-treated rats.. There were no group differences during the extinction retrieval test. Retrieval data from male animals are shown in grey and dark blue circles and squares, while white and light blue circles and squares represent data from female animals. **(C)** Schematic showing cannula placements in the IL. Data are expressed as means ± standard error of the mean (s.e.m.) [VEH-VEH (*n* = 13); VEH-PIC (*n* = 9); ANISO-VEH (*n* = 11); ANISO-PIC (*n* = 8)]; ANISO = anisomycin, BL = baseline, PIC = picrotoxin, VEH = vehicle.

As shown in Figure 4B (left panel), all groups showed similar increases in freezing levels during the conditioning session. This was confirmed by a three-way ANOVA, which showed a main effect of trial (*F*_5, 180_ = 40.3, *p* < 0.01), but no other main effects or interactions (*Fs* < 0.62, *p*s *>* 0.43). Immediately after fear conditioning, animals received intra-IL infusions of anisomycin (ANISO) or VEH, on the next day were treated with intra-IL injections of PIC or VEH and were extinguished two days later. As shown in Figure 4B (middle panel), all groups had similar levels of freezing during the baseline period of the extinction session (*Fs* < 0.42, *p*s > 0.51). However, once extinction commenced the VEH-PIC group exhibited lower levels of freezing than all other groups. This was confirmed in a three-way ANOVA that revealed a main effect of trial (*F*_*9, 324*_ *=* 19.2, *p* < 0.01) and significant interactions between the two drug treatments (*F*_*1, 36*_ *=* 5.11, *p* = 0.03), as well as a significant three-way interaction (*F*_*9, 324*_ = 2.85, *p* < 0.01). Similar to the first experiment, PIC did not affect freezing during the first CS (Figure S1).

The next day animals were returned to the extinction context for a retrieval test. As shown in Figure 4B (right panel), no differences were observed between groups. This was confirmed by a two-way ANOVA which showed no significant main effects or interactions (*F*s < 0.71, *p*s > 0.41). This contrasts with the outcome we observed in Experiment 2, in which PIC treated animals exhibited less freezing during the extinction retrieval test compared to controls. The reason for this difference is not clear, though it is noteworthy that the variability in the VEH-PIC group was very high in this experiment. Ultimately, these data suggest that protein synthesis in the IL during the consolidation of fear is required PIC for the suppression of conditioned freezing by IL picrotoxin infusions.

## DISCUSSION

In this study we show that IL picrotoxin infusions from minutes to days after auditory fear conditioning attenuate conditional freezing and facilitate fear extinction retrieval. Although the effects of pharmacological stimulation of the IL are not restricted to the classic consolidation time-window, protein synthesis inhibition in the IL immediately after fear conditioning attenuated the effects of picrotoxin infusions performed 24 h later. This suggests that protein synthesis in the IL soon after fear conditioning contributes to subsequent extinction learning. Interestingly, we found that fear memories are susceptible to IL picrotoxin up to 13 days after fear conditioning. This reveals that post-training picrotoxin modulates IL processes involved in extinction learning long after cellular consolidation of the fear memory is complete. Ultimately, the present study reveals that fear conditioning recruits the IL to enable subsequent fear extinction.

It is possible that the effects we observed were due to a disruption of fear memory consolidation by IL picrotoxin infusions. However, this is unlikely because freezing to the initial CS in each of the four experiments was spared. Indeed, early lesion studies indicate that the IL is not necessary for fear memory formation or retrieval [3,4]. Moreover, animals that received both anisomycin immediately after fear conditioning and picrotoxin 24 h later did not show a suppression of freezing. This suggests that picrotoxin did not disrupt fear memory consolidation, but rather augmented an inhibitory memory that was established shortly after fear conditioning. In all four experiments, this resulted in a reduction in conditioned freezing during the extinction training session, and resulted in lower conditioned freezing during the extinction retrieval test when PIC was delivered within 24 hours of conditioning (Experiments 1 and 2). Unexpectedly, enhanced extinction retrieval was not observed in Experiment 4 (in which intra-IL IC infusions also were made 24 hrs after conditioning). Methodological differences regarding the timing and number of drug manipulations may have masked this effect.

Several studies have demonstrated that IL manipulations performed during extinction training can bidirectionally modulate extinction retrieval the next day [4, 5, 9, 26-28]. Several lines of evidence suggest these effects rely on reciprocal connections between the IL and amygdala [20, 29-31] and are regulated by projections from the ventral hippocampus to the IL [32, 33]. In all of these cases, activity in the IL or its projections were manipulated during either the extinction learning or retrieval sessions. Here we show for the first time that IL activation up to 13 days before extinction learning has a similar effect to manipulations of IL activity during or after extinction learning. It will be of considerable interest to determine whether post-conditioning intra-IL PIC infusions suppress freezing and augment extinction learning by not only influencing local plasticity within the IL (which is suggested by Experiment 4), but also promoting plasticity in IL efferents associated with extinction learning such as amygdala intercalated cells [34, 35] or the nucleus reuniens [36, 37].

The finding that IL is engaged during the consolidation of fear conditioning to drive future extinction learning is paradoxical. However, recent studies have shown that disrupting IL activity up to 12 h after fear conditioning causes extinction deficits [14, 15]. Additionally, it was previously reported that intra-IL BDNF treatment performed 24 h after learning is sufficient to disrupt freezing, mimicking an extinction session [23]. We have now replicated this effect (Experiment 3) and shown that a similar outcome can be achieved with direct pharmacological stimulation of IL neurons. Collectively, this work suggests that cellular processes in the IL lasting hours to days after fear conditioning are fundamental for learning that occurs during extinction. Taken together, these findings show that the role for the IL in regulating conditioned freezing is not temporally restricted to the extinction learning experience.

This work points to the interesting possibility that the IL has a general role in inhibitory learning processes that occur in parallel with excitatory fear conditioning. This is consistent with reports that the IL is involved in a variety of inhibitory learning processes, including latent inhibition [12, 13] and conditioned inhibition [38]. Accordingly, we propose that post-conditioning infusions of either BDNF or picrotoxin augment the consolidation of an inhibitory engram in the IL that is formed during (or shortly after) fear conditioning. We propose that this inhibitory engram promotes the suppression of freezing during subsequent extinction training and retrieval tests. This process appears to involve protein synthesis in the IL insofar as anisomycin injected in the IL immediately after fear conditioning prevents the effects of postconditioning picrotoxin. This is consistent with recent work showing that social buffering of fear induces plasticity in the IL, and optogenetic reactivation of IL neurons that were active during social interaction is sufficient to suppress later freezing [39]. Inhibitory engrams may be mediated by plasticity at inhibitory synapses and have been postulated to underlie the suppression of behavioral responses that occurs during habituation, for example [40].

The establishment of excitatory and inhibitory engrams during fear conditioning has theoretical underpinnings in opponent-process theory [41]. In opponent process theory, a biologically significant event activates 1) an A-process that drives an immediate emotional response and 2) a temporally delayed B-process that opposes the emotional response associated with the A-process. Our results suggest that during fear conditioning, the IL encodes a B-process that opposes the aversive properties of the US. Although there are very few studies on the contribution of opponent processes to fear conditioning, Fanselow & Tighe (1988) suggested that an inhibitory opponent process defines the effective inter-trial interval in contextual conditioning [42]. Inhibitory engrams encoding opponent processes experienced during fear conditioning may have adaptive value by priming the network for future memory updating.

The observation that intra-IL picrotoxin or BDNF injections [43] facilitate extinction when injected two weeks after fear conditioning is intriguing and demands further investigation. Interestingly, Peters and colleagues (2010) found that when the BDNF injections were performed 24 h before conditioning, there was no effect of drug [23]. This indicates that whatever the mechanism supporting fear suppression, it depends on fear conditioning. Our current understanding of cellular memory consolidation does not explain how a memory becomes susceptible to pharmacological stimulation two weeks after it is acquired. One possibility is that the drug administration procedures (handling and transport) reactivate the fear memory and render it susceptible to disruption by impairing reconsolidation [44]. Further work is required to explore this possibility.

In sum, our work suggests that the IL encodes an inhibitory memory during the consolidation of excitatory auditory fear conditioning. This inhibitory trace opposes fear and is promotes the subsequent extinction of fear. Future studies employing engram-tagging techniques are necessary to understand the nature and function of the inhibitory engram encoded in the IL during fear memory consolidation.

## Author contributions

H.B. and S.M. designed all the experiments and analyses. H.B., J.H.Jr, H.L.V., G.M.G. and V.A.L.J. performed stereotactic surgeries. H.B., J.H.Jr, C.R.O., H.L.V., G.M.G and V.A.L.J. performed behavioral experiments. H.B. analyzed the data. H.B. and S.M. wrote the manuscript.

## SUPPLEMENTAL MATERIAL

### Picrotoxin spares conditioned freezing in the first extinction trial

In four experiments we found that picrotoxin infused from 1 hour to 13 days after fear conditioning produced dramatic impairments in conditioned freezing during subsequent extinction sessions. Freezing during the first 5-trial block was consistently reduced in each experiment. However, it is important to know whether freezing was lower in response to the first trial of each extinction session, because this trial provides an index of fear memory retrieval independent of extinction. To this end, we examined freezing during each of the first five conditioning trials for all the experiments. Each trial consisted of a 10-sec tone and a 30-sec inter-trial interval (ITI). We only compared animals that received vehicle or PICRO in the IL (BDNF- and ANISO-treated animals were excluded from this analysis).

As shown in Figure S1 (A), conditioned freezing was similar in VEH- and PICRO-treated rats during the first CS presentation of the extinction session but diverged during the first ITI in each of the four experiments. Figure S1 (B) shows freezing during the first CS in each experiment. In none of the experiments did the difference between VEH and PICRO groups reach significance (*p*’s > 0.053). These analyses suggest that even though the PICRO manipulations generally reduced freezing during the first 5-trial extinction block, the infusions spared freezing to the first CS of the extinction session. Picrotoxin infusions into the IL appears to accelerate extinction of freezing to the CS rather than producing a general reduction in the expression of the conditioned freezing response or retrieval of the fear memory.

**Figure S1.**
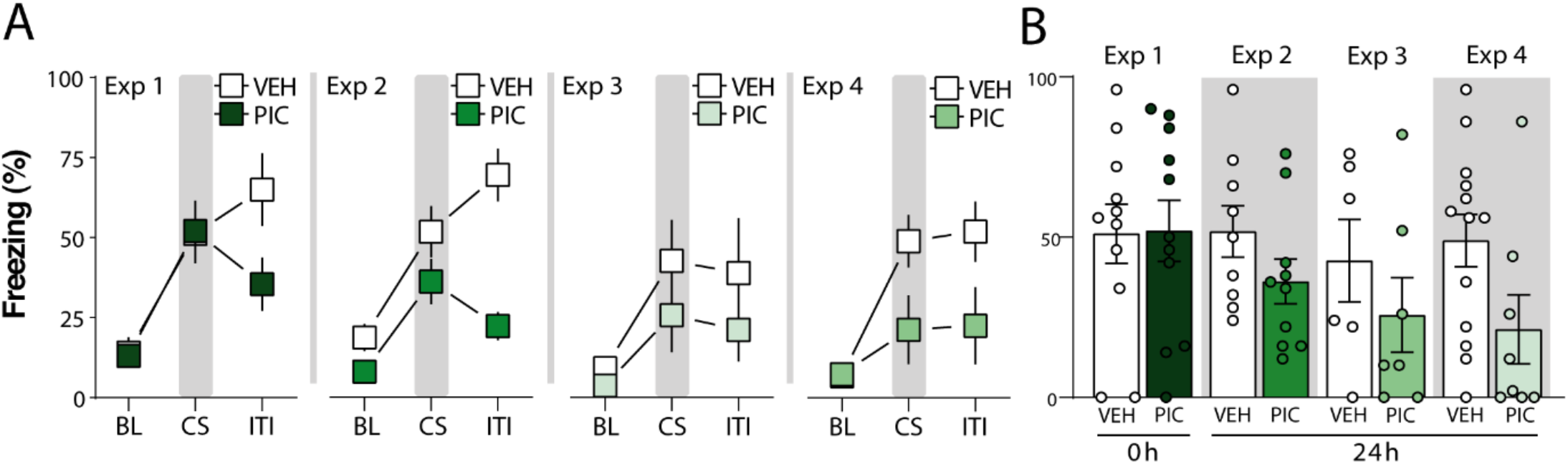
Picrotoxin does not disrupt fear recall on the first extinction trial. **(A)** Freezing data during the first extinction trial during the baseline (BL) period, the conditioned stimulus (CS), and post-CS intertrial intertrial (ITI) for each experiment. Freezing during the CS is highlighted by gray bars. **(B)** Average CS freezing in each of the four experiments in VEH- and PIC-treated rats. There were no group differences in any of the experiments (Exp 1: *p* = 0.94; Exp 2: *p* = 0.15; Exp 3: *p* = 0.35; Exp 4: *p* = 0.053). Exp = experiment; PIC = picrotoxin; VEH = vehicle.

